# A complex of the lipid transport ER proteins TMEM24 and C2CD2 with band 4.1 at cell-cell contacts

**DOI:** 10.1101/2023.12.06.570396

**Authors:** Ben Johnson, Maria Iuliano, TuKiet Lam, Thomas Biederer, Pietro De Camilli

## Abstract

Junctions between the ER and the plasma membrane (ER/PM junctions) are implicated in calcium homeostasis, non-vesicular lipid transfer and other cellular functions. Two ER proteins that function both as membrane tethers to the PM via a polybasic motif in their C-terminus and as phospholipid transporters are brain-enriched TMEM24 (C2CD2L) and its paralog C2CD2. Based on an unbiased proximity ligation analysis, we found that both proteins can also form a complex with band 4.1 family members, which in turn can bind a variety of plasma membrane proteins including cell adhesion molecules such as SynCAM 1. This complex results in the enrichment of TMEM24 and C2CD2 containing ER/PM junctions at sites of cell contacts. Dynamic properties of TMEM24-dependent ER/PM contacts are impacted when in complex as TMEM24 present at cell adjacent junctions is not shed by calcium rise, unlike TMEM24 at non-cell adjacent junctions. These findings suggest that cell-contact interactions control ER/PM junctions via TMEM24 complexes involving band 4.1 proteins.

**SUMMARY:** The ER-anchored lipid transfer proteins TMEM24/C2CD2L and its paralog C2CD2 mediate formation of ER-plasma membrane junctions at sites of cell-cell contacts by interacting with band 4.1 family members and indirectly with cell adhesion proteins.

## INTRODUCTION

Junctions between the endoplasmic reticulum and the plasma membrane (ER/PM junctions) are sites where the ER is held in close apposition (∼ 15-25 nm distance) to the PM by protein tethers without undergoing fusion (Chung et al., 2022; Elbaz and Schuldiner, 2011; Fernández-Busnadiego et al., 2015; Rosenbluth, 1962; Wu et al., 2018). Most ER/PM tethers have additional roles beyond holding the two membranes together (Balla et al., 2020; Chen et al., 2019; Gatta and Levine, 2017; Johnson et al., 2018; Kirmiz et al., 2018; Saheki and De Camilli, 2017; Scorrano et al., 2019; Wu et al., 2018). The abundance of ER/PM junctions varies from cell type to cell type and, within a given cell type, they can be heterogenous in molecular composition and morphology, such as area of membrane apposition and width of the ER lumen. Their abundance and structure can also differ on PM surfaces that face different neighbors (Chang et al., 2017; Chung et al., 2022; Deardorff et al., 2014; Wu et al., 2017). Moreover, ER/PM junctions can be populated by different proteins and undergo expansion and reduction in size depending upon the functional state of the cell (Chang et al., 2017; Giordano et al., 2013). In neurons, where ER/PM junctions can cover as much as 10% of the PM (Kuijpers et al., 2023; Wu et al., 2017) such junctions can decrease in both size and number by ∼50% during depolarization with high K^+^ or NMDA treatment with differences in sensitivity being linked to junction morphology (Tao-Cheng, 2018).

Proteins that both localize at ER/PM junctions and also participate in their formation include ion channels such as Kv2 potassium channels in neurons (Fox et al., 2015; Johnson et al., 2018; Kirmiz et al., 2018), the ER protein STIM1, which plays a critical role in the regulation of store-operated Ca^2+^ entry via its binding the PM Ca^2+^ channel Orai (Chang et al., 2017), as well as a multiplicity of proteins implicated in the non-vesicular transport of lipids (Amos et al., 2023; Chang et al., 2017; Chang and Liou, 2013; Chung et al., 2023; Johnson et al., 2019; Saheki and De Camilli, 2017; Suzuki et al., 2014; Thakur et al., 2019; Yeun et al., 2015). This mode of lipid transport is critically important as most lipid species in eukaryotic cells are synthesized within the ER. Thus, homeostasis of PM lipids, which is needed to support the structural and signaling functions of this membrane, requires rapid and efficient exchanges of lipids between the ER and the PM both for the delivery of newly synthesized lipids and for the return to the ER of lipid catabolites for metabolic recycling (Guillén-Samander and De Camilli, 2023).

A protein that functions both as an ER/PM tether and as a lipid transporter is TMEM24 (also called C2CD2L), which is primarily expressed in neurons and pancreatic β-cells (Lees et al., 2017; Sun et al., 2019; Xie et al., 2022). TMEM24 is anchored into the ER membrane via an N-terminal transmembrane region, which is followed on the cytosolic side by an SMP domain, a C2 domain, and an approximately 300 amino acid long region which is predicted to be primarily unstructured, but contain stretches of high conservation, including a polybasic C-terminal sequence (Lees et al., 2017; Sun et al., 2019). TMEM24 dimerizes via the SMP domain, which is the lipid transport module and harbors glycerolipids. It binds the PM through a charge-based association between its polybasic motif and negatively charged lipids present on the inner leaflet of the PM. This interaction is negatively regulated by the Ca^2+^ and PKC-dependent phosphorylation of serine residues interspersed within the polybasic motif. Accordingly, TMEM24 expressed in neurons or other cells is reversibly released from ER/PM junctions and redistributes throughout the ER in response to cytosolic Ca^2+^ elevations (Lees et al., 2017; Sun et al., 2019; Xie et al., 2022). TMEM24 has a paralog, CDCD2, which in contrast to TMEM24 is broadly expressed in all tissues. While having the same domain structure as TMEM24, C2CD2 lacks the serine residues responsible for the phosphoswitch that controls the PM tethering function of TMEM24, and thus is not released from ER/PM junctions in response to Ca^2+^ elevations (Sun et al., 2019).

While studying the properties of TMEM24 in semiconfluent cultured cell lines we observed a non-homogenous distribution of TMEM24 between portions of the PM adjacent, or non-adjacent, to other cells. Here we have elucidated the molecular mechanisms responsible for this heterogenous localization of TMEM24 and have discovered an interaction of TMEM24 with band 4.1 proteins, which in turn link TMEM24, and its paralogue C2CD2, to cell adhesion molecules. We speculate that the lipid transport function of TMEM24 may play a role in the support of the signaling reactions that occur at these sites.

## RESULTS

### Preferential accumulation of TMEM24 & C2CD2 at ER/PM junctions localized at sites of cell-cell contact

When TMEM24-eGFP or its paralog C2CD2-eGFP were expressed in HEK293 cells they localized at patches along the PM with the expected pattern of ER/PM junctions. However, such patches were significantly larger at sites of cell-cell contact (Figure 1A). Often these larger junctions spanned the entirety of the cell-cell interface in our overexpression system which is expected to result in expansion of endogenous ER/PM contacts. The preferential accumulation to sites where the PMs of two cells are in close apposition was unique to TMEM24 and C2CD2. Other ER/PM tethering proteins that we tested, such as Junctophilin-4 (JPH4-eGFP), extended synaptotagmin-2 (eGFP-E-Syt2), Kv2.1-GFP, or the constitutively active YFP-STIM1(D76A) mutant, showed no difference in size based on the presence of an adjacent cell (Figure 1A and Supplemental Figure 1, quantified in Figure 1B).

**Figure 1.**
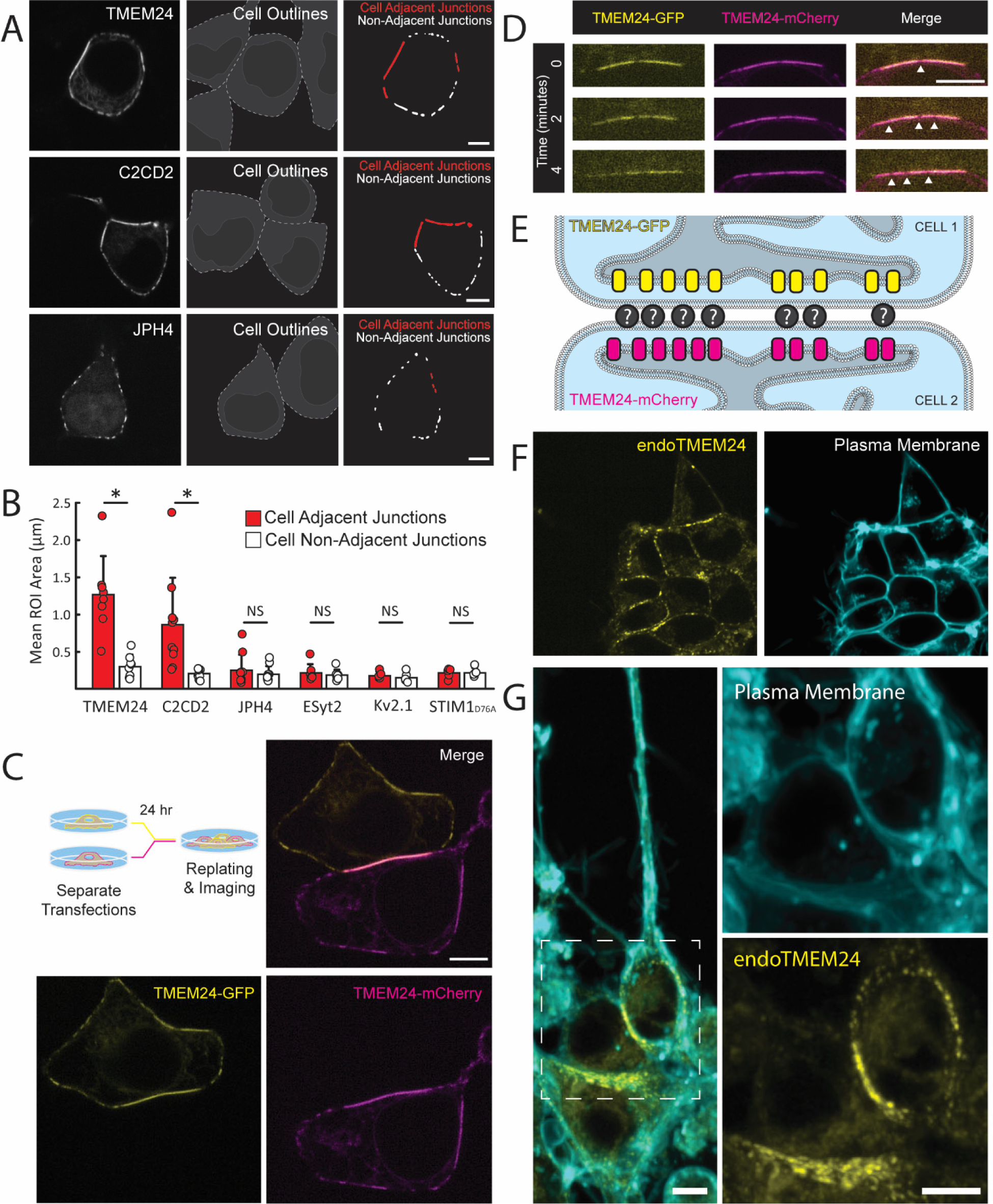
Large TMEM24-positive ER/PM junctions at cell-cell interfaces. **(A)** Left, Spinning disk confocal images of TMEM24-mCherry, C2CD2-eGFP, and JPH4-eGFP in HEK293 cells. Center, cell silhouettes obtained by cell labeling with Tracker DiI or CellTracker Red. Right, ER/PM junctions from the left fields with cell-adjacent ER/PM junctions outlined in red. **(B)** Quantification of ER/PM junction sizes at cell adjacent and non-adjacent regions of the PM. Student’s t-test p-values of *p* = 0.0096 for TMEM24 and *p* = 0.01 for C2CD2. **(C)** and **(D)** Cells expressing TMEM24-eGFP and TMEM24-mCherry were co-plated as illustrated by the diagram. **(C)** TMEM24 expressed in adjacent cells forms symmetrical ER/PM junctions. **(D)** Gaps in the TMEM24 signal (indicated by arrowheads) in one cell are mirrored by gaps in the signal of the adjacent cell. **(E)** Diagram illustrating TMEM24 behavior in adjacent cells expressing TMEM24. **(F)** TMEM24 tagged with GFP at the endogenous locus in IMR32 cells localizes to the cell-cell interface. **(G)** Differentiated IMR32 cells with TMEM24 tagged at the endogenous locus also display a similar localization pattern. Plasma membrane labeled with CellBrite 650 dye in **(F)** and **(G)**. Scale bars = 5 μm.

We next expressed TMEM24-eGFP in one set of HEK293 cells, TMEM24-mCherry in another set of HEK293 cells, and after 24 hours co-plated the two cell populations together (Figure 1C). This protocol allowed us to observe the dynamics of TMEM24-positive ER/PM junctions across neighboring cells. Cell pairs that expressed TMEM24 in both cells formed large symmetrical ER/PM junctions across the cell-cell interface with gaps that appeared in the TMEM24 signal over time of one cell being perfectly mirrored by simultaneous gaps in the signal of the neighboring cell (Figure 1D). In contrast, expression of tagged TMEM24 in one cell and either tagged JPH4 or E-Syt2 in an adjacent cell did not result in such large symmetrical junctions (Supplemental Figure 2). Cell pairs expressing other ER/PM tethers (eGFP-E-Syt2 & mCh-E-Syt2, JPH4-eGFP & JPH4-mCh) also did not generate these robustly expanded symmetrical junctions, though expression of JPH4 in adjacent cells generated perhaps slight symmetry which we did not further explore (Supplemental Figure 2). These data strongly suggest that TMEM24 may be part of a direct or indirect complex with endogenously expressed cell adhesion molecule(s) (see Figure 1E for model).

To ensure that the localization of TMEM24 was not a result of overexpression, we also analyzed the localization of endogenous TMEM24. To this aim we used previously generated IMR32 human neuroblastoma cells in which endogenous TMEM24 (endoTMEM24-eGFP) had been tagged at the TMEM24 gene locus by gene editing (Sun et al., 2019). In agreement with the HEK293 cell data, we found that also in IMR32 cells TMEM24 accumulated at ER/PM junctions localized at sites of cell appositions both before and after brdU-induced neuronal differentiation (see Figures 1F&G). This demonstrates that the localization of TMEM24 at sites of cell adhesion is not an artifact of an overexpression system and that it is also not unique to HEK293 cells.

### TMEM24 cell-adjacent and cell non-adjacent ER/PM junctions have distinct properties

Co-expressing TMEM24 with other ER/PM tethers revealed molecular heterogeneity between cell-adjacent and cell non-adjacent ER/PM junctions. JPH4-eGFP and eGFP-E-Syt2 colocalized with TMEM24-mCherry at ER/PM junctions that are cell non-adjacent but were excluded from TMEM24-mCherry-positive ER/PM junctions localized at sites of cell-cell interfaces (Figure 2A&B and Supplemental Fig. 2). In contrast, Kv2.1-GFP and YFP-STIM1(D76A) colocalized with TMEM24-mCh at both cell-adjacent and cell non-adjacent junctions (Supplemental Figure 2).

**Figure 2.**
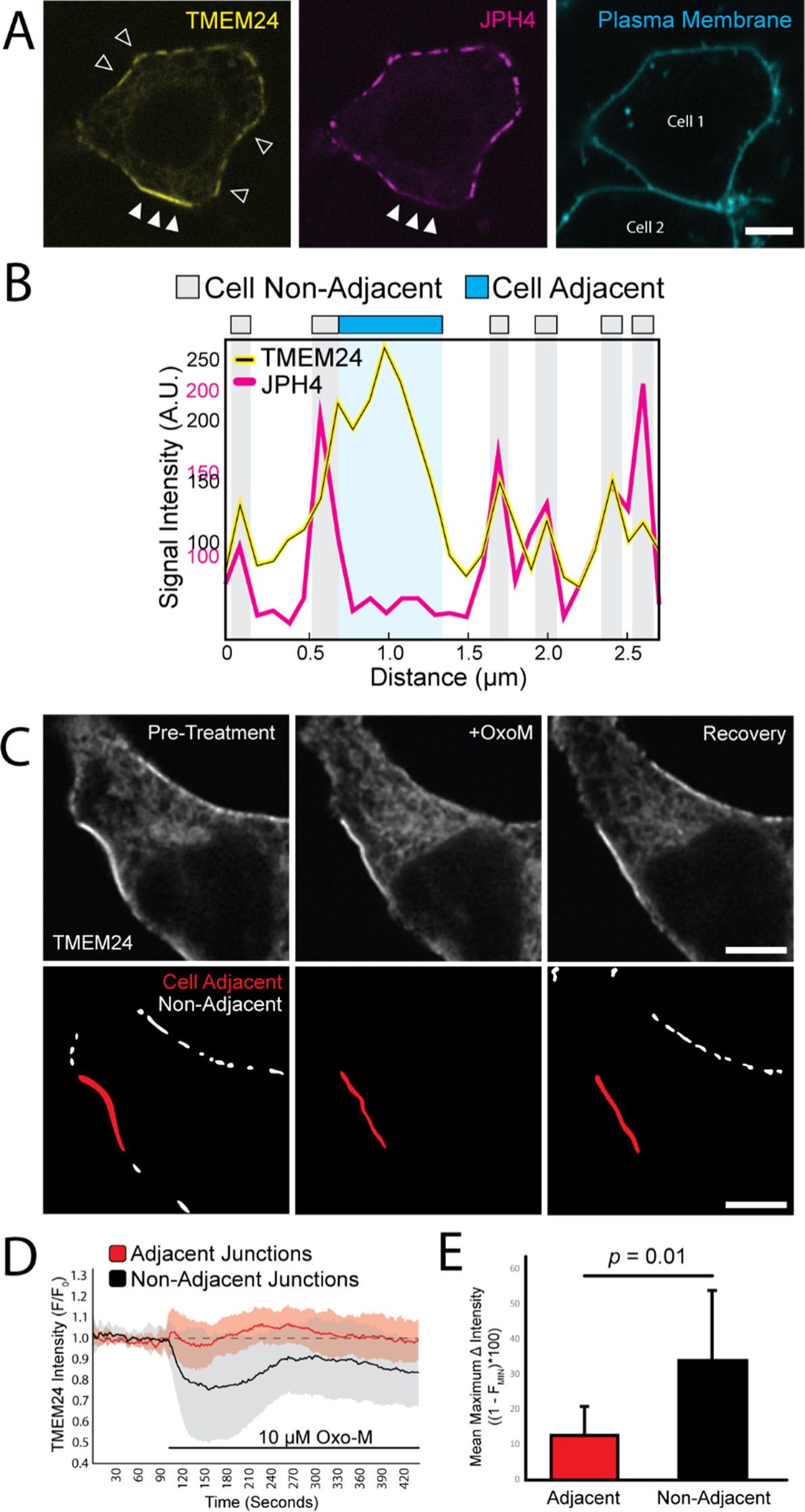
TMEM24-positive cell-adjacent junctions exhibit distinct characteristics. **(A)** TMEM24-mCherry and JPH4-eGFP colocalize at ER/PM junctions that are non-cell adjacent (open arrowheads represent a few examples contained within the image) but JPH4 is excluded from TMEM24-induced cell-adjacent ER/PM junctions (solid arrowheads). Plasma membrane labeled with CellBrite 650. **(B)** Representative line scan of a cell membrane demonstrating fluorescence increases in both TMEM24 and JPH4 channels at ER/PM junctions with the exception of the cell-cell contact area where TMEM24 signal increases and JPH4 signal is lacking. **(C)** Images of TMEM24-mCherry cell adjacent and cell non-adjacent ER/PM junctions before and after addition of 10 μM Oxotremorine-M (OxoM). Junctions that are cell-adjacent and cell non-adjacent are indicated in lower panels. **(D)** TMEM24 response to treatment of 10 μM OxoM separated by cell-adjacent and cell non-adjacent TMEM24 fluorescence over time and normalized to an average pre-treatment fluorescence level. **(E)** Maximal average fluorescence change in TMEM24 signal at cell-adjacent and cell non-adjacent ER/PM junctions. Student’s t-test returned a value of *p* = 0.01. All images are from HEK293 cells and were acquired by confocal microscopy. Scale bars = 5 μm.

We previously showed that the localization of TMEM24 at the PM is regulated by cytosolic calcium (Lees et al., 2017; Sun et al., 2019). Calcium/PKC-dependent phosphorylation of basic amino acid residues within the C-terminal polybasic motif of TMEM24 disrupts its binding to the cytosolic surface of the PM (which is enriched in negatively charged phospholipids) and induces its redistribution throughout the ER. Conversely, dephosphorylation of the same residues by the phosphatase calcineurin/PP2B results in the rapid reassociation of TMEM24 with the PM. While TMEM24 was rapidly shed from the PM at cell non-adjacent ER/PM junctions in response to cytosolic calcium elevation produced by Oxo-M-induced M1 receptor activation, the localization of TMEM24 at junctions that are cell-adjacent was resistant to this disruption (Figures 2 C-E). This demonstrated that TMEM24 pools at cell-adjacent and cell non-adjacent junctions responded differently to external stimuli.

### A *β*-Sheet within the TMEM24 C-terminal region is necessary and sufficient for localization at sites of cell-cell adhesion

Truncation and internal deletion mutations of TMEM24 were generated to identify the region(s) responsible for its localization at sites of cell-cell contacts (see schematic domain representation of TMEM24 in Figure 3A) A construct comprising amino acids 1-414 and including the ER transmembrane sequence and the SMP and C2 domains was localized throughout the ER and failed to accumulate at any ER/PM junction (Figure 3B), while a longer construct (a.a. 1-630) accumulated at ER/PM junctions, but selectively at sites of cell-cell contact (Figure 3C). As the 1-630 construct lacks the phospho-regulated PBM (a.a. 666-706) previously shown to interact with the acidic leaflet of the PM bilayer (Lees et al., 2017; Sun et al., 2019), this finding suggested that TMEM24 contained an additional PM tethering motif within its 414-630 region allowing it to form junctions specifically at cell-cell interfaces. To confirm this hypothesis, we generated a hybrid construct comprising the TMEM24 414-630 region fused to the ER protein TRAPγ-eGFP. TRAPγ is a component of the translocon complex that localizes diffusely and homogenously throughout the ER when expressed alone (Figure 3D). The fusion protein, TRAPγ-TMEM24(414-630)-eGFP, not only accumulated at ER/PM junctions but did so specifically at sites of cell-cell contact (Figure 3E). AlphaFold predictions suggest that the 414-630 region of TMEM24 is predominantly disordered, but contains three short, structured a.a. sequences: two ⍺-helices and one β-sheet (Figure 3F) (Jumper et al., 2021). An internal deletion of the first three strands of the β-sheet, in combination with phosphomimetic mutations of the 5 serine residues (S→E) shown to abolish the interaction of the PBM of TMEM24 with the acidic cytosolic leaflet of the PM (Sun et al., 2019), completely disrupted TMEM24’s ability to bind the PM, suggesting that the β-sheet is necessary for its localization at sites of cell-cell contacts (Figure 3G). AlphaFold predicts a similar β-sheet in the corresponding region of C2CD2. These regions of both TMEM24 and C2CD2 are highly conserved across higher organisms (Supplemental Figure 3).

**Figure 3.**
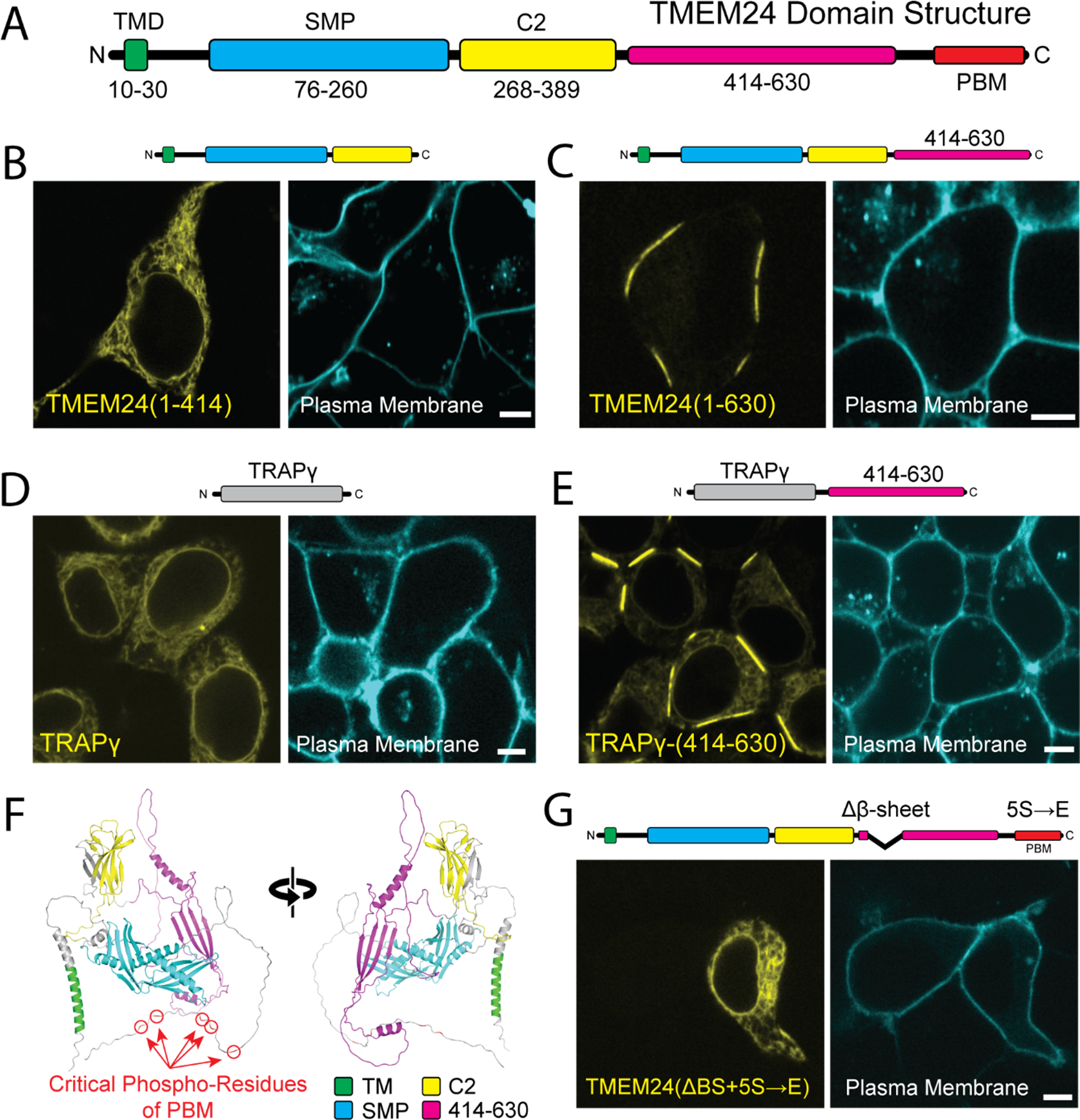
A *β*-sheet within the TMEM24 C-terminus is responsible for its targeting to cell-adjacent ER/PM junctions. HEK293 cells. **(A)** TMEM24 domain architecture. TMR = transmembrane region; PBM = polybasic motif. **(B)** Fragment 1-414 is diffusely ER localized and does not localize to ER/PM junctions at any location. **(C)** Fragment 1-630 is localized at ER/PM junctions at sites of cell-cell contact **(D)** The TRAPγ-eGFP has a diffuse ER localization. **(E)** TRAPγ-(414-630)-eGFP hybrid construct localizes to ER/PM junctions at sites of cell-cell contact. **(F)** Alphafold predicted structure of a TMEM24 monomer in two different orientations. **(G)** Disruption of both the polybasic motif (accomplished via the 5S→E mutation) and the TMEM24 C-terminal β-sheet interferes with TMEM24’s ability to localize at any ER/PM junction. Plasma membranes were labeled with CellBrite 650. Scale bars = 5 μm.

### Identification of TMEM24 interactors via APEX2 proximity biotinylation

To identify potential PM protein interactors for TMEM24 that may be responsible for its targeting to sites of cell adhesion, we turned to APEX2-based proximity biotinylation. In the presence of H_2_O_2_, APEX2-conjugated proteins generate rapidly diffusing biotin radicals that biotinylate electron-rich amino acid residues of nearby proteins. As these radicals have extremely short half-lives (<1 ms), only proteins in the immediate vicinity (∼20 nm) of the APEX2 protein during H_2_O_2_ treatment are biotinylated. Biotinylated proteins can be subsequently purified via streptavidin-based affinity purification and identified by mass spectrometry. This approach has already been successfully used for interrogating the proteome of the ER/PM junction (Jing et al., 2015; Johnson et al., 2018) as well as other cellular subcompartments such as the synaptic cleft (Cijsouw et al., 2018; Loh et al., 2016).

Full length TMEM24 fused to APEX2 and several other control constructs were generated (Figure 4A). All these constructs were additionally tagged with eGFP to validate their correct targeting by fluorescence microscopy (Figure 4A). eGFP-APEX2-Sec61β and TMEM24(1-414)-APEX2-eGFP provided controls for diffuse ER localization. TMEM24(1-630)-APEX2-eGFP and TMEM24(1-666)-APEX2-eGFP, which both preferentially localized at cell-adjacent ER/PM junctions, also served as reporters. Untransfected HEK293 cells were used to discriminate between the signal produced by the transfected constructs and endogenous biotinylation. After the APEX2 reaction, a pool of cells were fixed and labeled with a streptavidin-conjugated far-red probe. Microscopy analysis of these cells confirmed that cell regions intensely positive for the biotinylation signal were consistent with the subcellular localization of each eGFP-labeled construct (Figure 4A). Another pool of cells was homogenized, biotinylated proteins were affinity-purified on streptavidin beads, and the affinity purified material analyzed by SDS-PAGE. Streptavidin and anti-GFP immunolabeling of this material revealed not only strong self-biotinylation of the APEX2-constructs as expected (identified by anti-GFP Western blotting), but also a wide range of additional biotinylated protein bands (Supplemental Figure 4). To detect the nature of these other proteins, affinity purified samples were submitted for mass spectrometry analysis in triplicate (three independent biological replicates).

**Figure 4.**
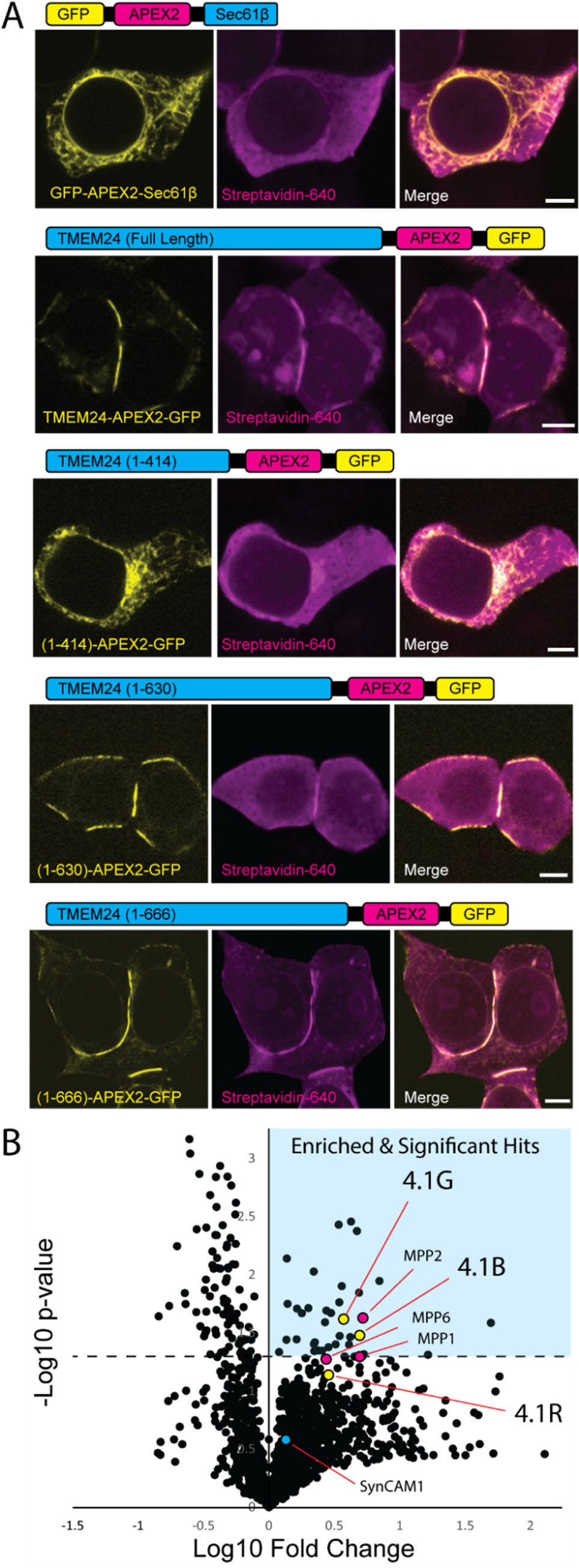
APEX2 proximity biotinylation identifies TMEM24 protein neighbors. **(A)** Schematic representations and confocal images of APEX2 conjugated proteins expressed in HEK293 cells and used in the APEX screen. After the APEX reaction cells were fixed, labeled with CF640R streptavidin to visualize biotinylated proteins and imaged via confocal microscopy. **(B)** Volcano plot of proteins identified via mass spectrometry after streptavidin-based purification. Significantly enriched proteins are found in the upper right quadrant indicated by cyan. Student t-test values for 4.1G, 4.1B, and 4.1R were found to be *p* = 0.024, *p* = 0.033, and *p* = 0.074, respectively. Scale bars = 5 μm

The lists of proteins identified by mass spectrometry in the different sets of affinity-purified biotinylated material were compared to identify hits specifically enriched in the samples biotinylated by full length TMEM24-APEX2-eGFP, TMEM24(1-630)-APEX2-eGFP and TMEM24(1-666)-APEX2-eGFP constructs, relative to controls lacking the a.a. 414-630 region (see Figure 4C for volcano plot of grouped data and Supplemental Table 1 for full list of identified proteins). Among the 36 significant hits (see Methods) there were two of the four band 4.1 proteins (4.1G and 4.1B) encoded by the human genome, which ranked at position 18 and 21 (*p* = 0.024, *p* = 0.033), respectively. A third band 4.1 protein, 4.1R, was also among the proteins enriched in constructs containing the a.a. 414-630 region (it ranked at position 64) although its enrichment did not reach significance (*p* = 0.074). The fourth band 4.1 family member, 4.1N, was not identified in any of our experimental or control APEX conditions and we did not test for endogenous expression in our HEK293 cells. Interestingly, TMEM24 was previously found as a potential interactor of 4.1R using a rat kidney yeast two-hybrid screen (Calinisan et al., 2005), although this interaction was not further validated. We note that the list of 36 significant hits includes desmoglien-2 (DSG-2), a protein that was recently shown to be part of symmetrical desmosome-endoplasmic reticulum subcellular complexes at cell-cell contacts (Bharathan et al., 2023).

### Band 4.1 proteins interact with the β-sheet of TMEM24 via their C-terminal domain

Band 4.1 family proteins, which lack a transmembrane region, are key components of the PM-associated cytoskeleton in all cells (Baines et al., 2014). They have a similar domain architecture (see Figure 5A, left) with the highest degree of conservation occurring within their N-terminal FERM domains (<u>F</u>our point 1, <u>E</u>zrin, <u>R</u>adixin, <u>M</u>oesin) and C-Terminal Domains (CTD) (Baines et al., 2014). The FERM domain mediates binding to the PM by interacting with a variety of PM localized proteins and lipids, including cell adhesion proteins such as SynCAM 1/CADM1, CADM4, CD44, and members of the β integrin family (Baines et al., 2014). The enrichment of three band 4.1 family members in the proximity proteome of TMEM24 constructs that accumulate at cell adhesion sites, along with their reported adaptor function for proteins implicated in cell adhesion, prompted us to explore a potential direct interaction of TMEM24 with members of the protein band 4.1 family.

**Figure 5.**
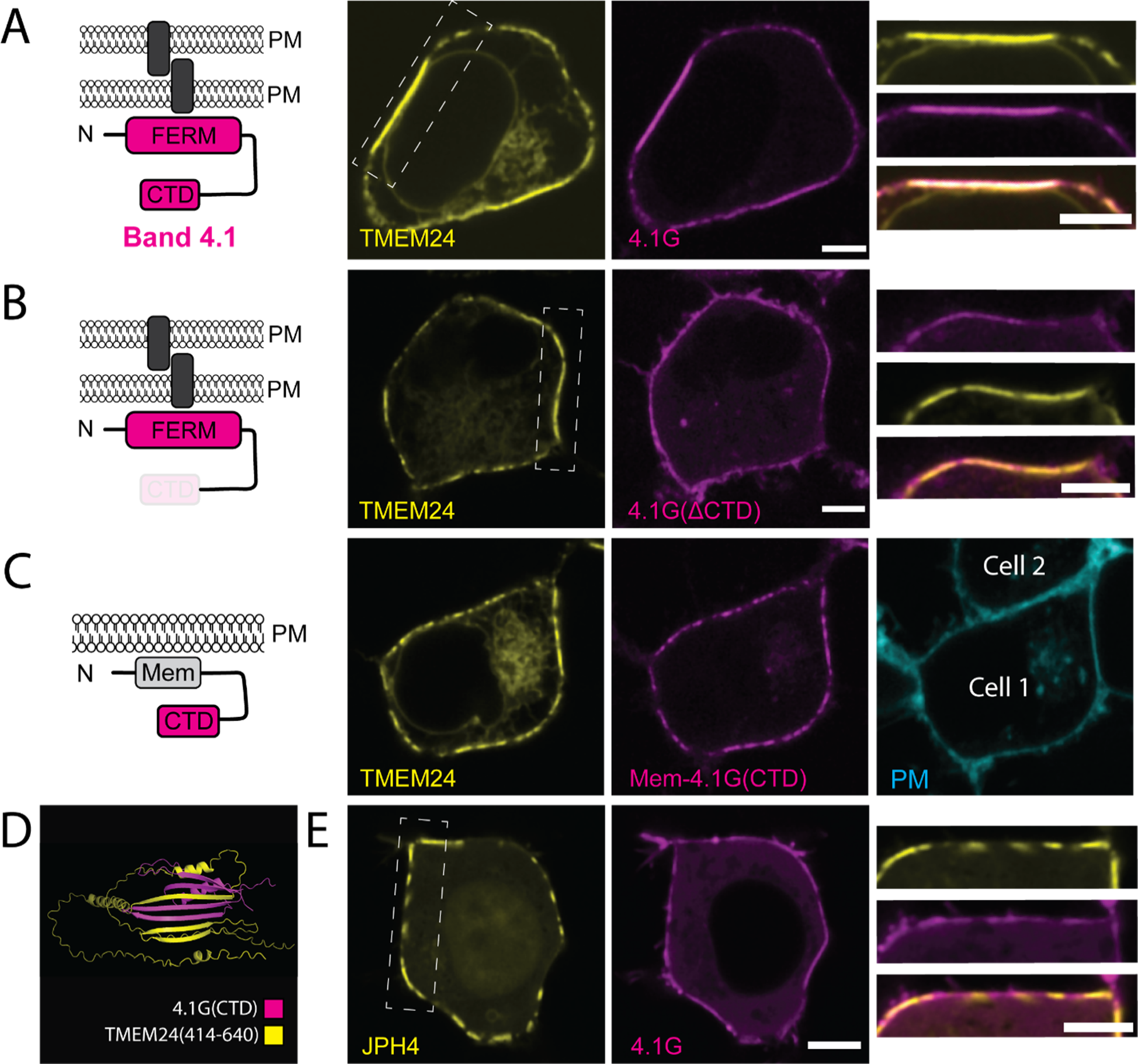
Band 4.1 Proteins bind TMEM24 via an interaction between TMEM24 *β*-sheet motif and Band 4.1 C-terminal domain. **(A)** Diagram of Band 4.1 family proteins (a cell adhesion protein is indicated by a gray rectangle) and confocal images of the indicated proteins coexpressed in HEK293 cells showing their colocalization. High magnification of the boxed regions of the middle panels are shown at right. **(B)** Diagram of Band 4.1 construct lacking the CTD and confocal images of the corresponding construct co-expressed with TMEM24 in HEK293 cells, showing lack of colocalization of the two proteins. **(C)** Diagram of the Mem-mCherry-4.1G(CTD) construct and confocal images of the corresponding construct co-expressed with TMEM24 in HEK293 cells. The two proteins are colocalized at ER/PM junctions, but the strong accumulation of TMEM24 at sites of cell-cell contacts is no longer observed. The panel at right shows the location of a neighboring cells stained with CellBrite 650. **(D)** Alphafold multimer predicted interaction between the β-sheet of TMEM24 and the β-sheet contained in the 4.1G CTD. **(E)** Coexpression of JPH4-eGFP and mCherry-4.1G in HEK293 cells result in no colocalization. Scale bars = 5 μm

To this aim, we co-expressed either mCherry-4.1G or mCherry-4.1R with TMEM24-eGFP in HEK293 cells (see Figure 5A and Supplemental Figure 5) and found that both proteins precisely colocalized at TMEM24 positive ER/PM contacts. The same results were obtained by co-expressing mCherry-4.1G and C2CD2-eGFP (Supplemental Figure 5). In contrast, a 4.1G construct that lacks its CTD (mCh-4.1GΔCTD) did not colocalize with TMEM24 (Figure 5B). Moreover, artificially targeting the CTD of 4.1G to the PM using a PM targeting peptide followed by a short linker (Mem-mCh-4.1(CTD)) was sufficient for colocalization with TMEM24 (Figures 5C). With this construct, however, which lacks the FERM domain, TMEM24 no longer became concentrated at sites of cell adhesion. These results indicate that TMEM24 binds the CTD of band 4.1 and that the FERM domain of band 4.1 is in turn responsible for targeting TMEM24 to sites of cell-cell contact.

As our studies described above had implicated the β-sheet motif within the C-terminal region of TMEM24 in its targeting to sites of cell-cell contact, we used Alphafold multimer (Evans et al., 2021), an algorithm designed to interrogate protein-protein interactions, to explore the presence in band 4.1 proteins of potential binding interfaces for this motif. This algorithm predicts with strong confidence (ranking confidence score of 0.75) the binding of the β-sheet of TMEM24 to the CTD of 4.1G, which also folds into a small β-sheet, with an intertwining of the β-strands of the two proteins to generate a single chimeric β-sheet (Figure 5D). Finally, when mCh-4.1G was coexpressed with another ER/PM tethering protein, JPH4-eGFP, it did not co-enrich with JPH4-positive ER/PM junctions and in fact appeared to be excluded from these sites (Figure 5E). Collectively, these results strongly suggest that the band 4.1 proteins function as adaptors between TMEM24 (via their interactions with its β-sheet motif) and cell adhesion molecules, via FERM-domain-dependent interactions.

### TMEM24 can form a complex with band 4.1 proteins and SynCAM 1

We next tested directly the possibility that TMEM24 and band 4.1 proteins could form a complex comprising also a cell adhesion protein. One cell-cell adhesion protein that binds the FERM domain of band 4.1 proteins is SynCAM 1 (also known as CADM1, Necl-2, TSLC-1, and IgSF4) (Baines et al., 2014; Yageta et al., 2002). We chose this protein to assess this possibility as endogenous SynCAM 1 was also a hit in our APEX screen in HEK293 cells, although its average enrichment relative to controls did not reach statistical significance. Moreover, SynCAM 1 in HEK293 cells was reported to form a tripartite complex comprising both band 4.1 proteins and members of the “membrane palmitoylated protein” (MPP) family (MPP1, MPP2 and MPP3) (Sakurai-Yageta et al., 2009) at cell-cell interfaces. Interestingly, 3 members of this family of MAGUK domain containing adaptors (MPP1, MPP2 and MPP6) were also hits in our screen for TMEM24 neighbors at sites of cell adhesion (see Figure 4B and Supplemental Table 1), with MPP1 being a statistically significant hit (ranked at position 16; *p* = 0.024). MPP2 and MPP6 were ranked at position 43 and 37 with p values of *p* = 0.053 and *p* = 0.0502, respectively.

To explore the ability of SynCAM 1 to form a complex comprising TMEM24 at sites of cell adhesion, we expressed SynCAM 1 with an extracellular eGFP tag at amino acid 363 (SynCAM 1(363)-eGFP) in HEK293 cells. This tag leaves the cytosolic sequence of SynCAM 1 unencumbered and also does not interfere with cell adhesion properties of the molecule (Fogel et al., 2007). When expressed alone, SynCAM 1(363)-eGFP was strongly enriched at cell-cell interfaces, although it was also present throughout the PM (note for example its uniform distribution on the basal surface of the cell), possibly due to overexpression (Figure 6A). However, when co-expressed with TMEM24-mCh, it co-clustered with this protein (most likely via the adapter role of endogenous 4.1 proteins) throughout the PM, both at the cell-cell interface as well as at all other ER/PM junctions including those formed at the basal surface (Figure 6B). In contrast, control co-expression of JPH4-mCh and SynCAM 1(363)-eGFP led to the exclusion of SynCAM 1 from JPH4-positive ER/PM junctions (Figure 6C), consistent with the exclusion of 4.1G from such junctions (Figure 5E) and the predominant exclusion of JPH4 from cell-adjacent junctions (Figure 2A & B). Finally, when expressed together, TMEM24, 4.1G, and SynCAM 1 colocalized as would be expected if they formed a complex (Figure 6D & E). These findings support the hypothesis that band 4.1 can function as an adaptor enriching TMEM24 at sites of cell adhesion.

**Figure 6.**
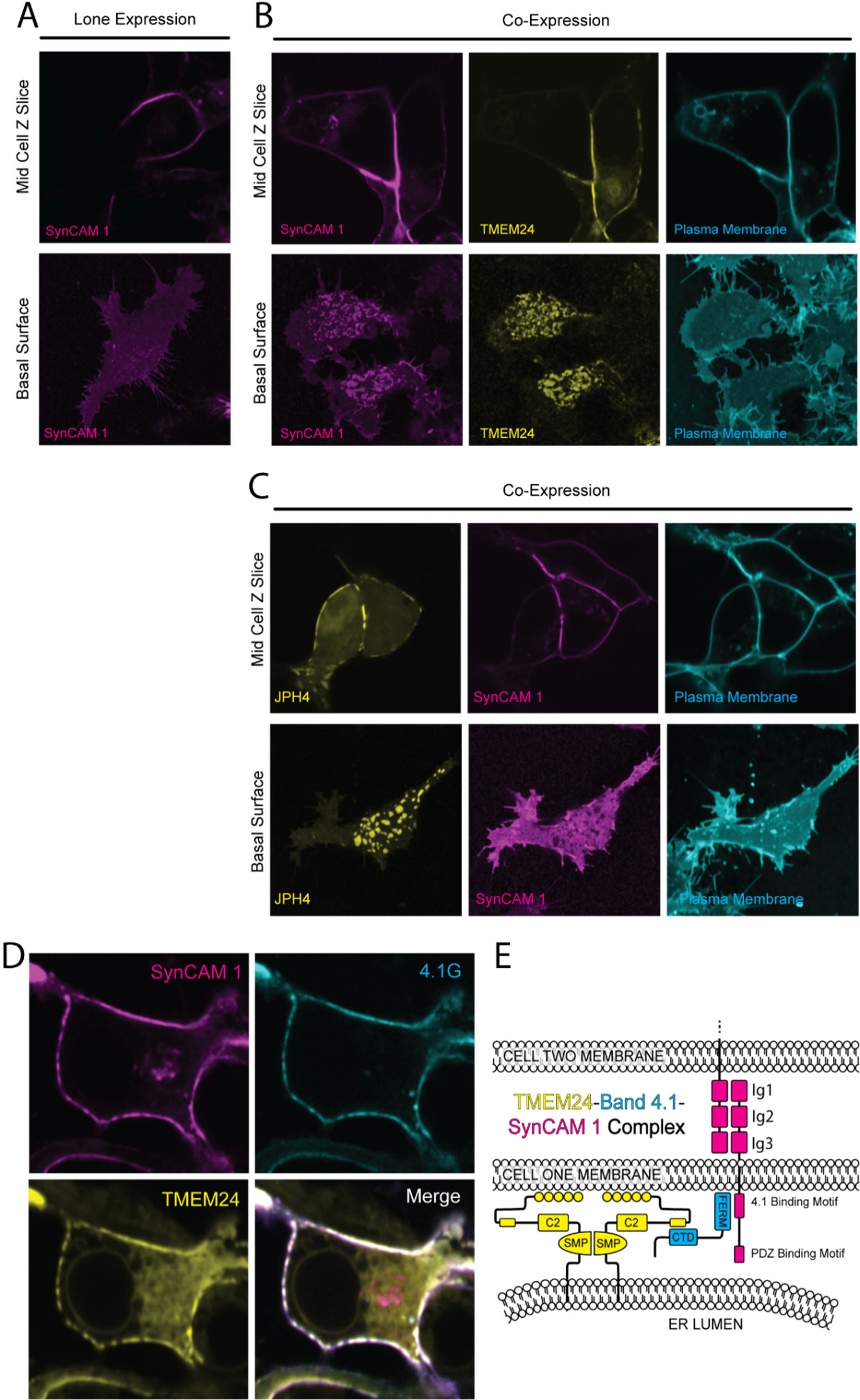
A TMEM24-4.1-SynCAM1 complex at sites of cell-cell contact. **(A)** Confocal images of SynCAM1(363)-eGFP expressed in HEK293 cells. The upper panel is a slice at mid z level while the lower panel is shows the basal surface. **(B)** Coexpression of TMEM24-mCherry and SynCAM1(363)-eGFP in HEK293 cells. Images are of a mid-level z-slice (above) or the basal surface (below) of the cell. Plasma membrane labeled with CellBrite 650. **(C)** Coexpression of JPH4-mCherry and SynCAM1(563)-eGFP in HEK293 cells. The plasma membrane was labeled with CellBrite 650. **(D)** Expression of SynCAM1(363)-eGFP, mCherry-4.1G, and TMEM24-Halo labeled with JF646 HaloTag ligand in HEK293 cells. All three proteins colocalize at ER/PM junctions. **(E)** Model of the TMEM24-4.1-SynCAM 1 complex. Scale bars = 5 μm

### TMEM24 ER/PM junctions can be found at neuronal contact sites

SynCAM 1 has been extensively studied in the nervous system. In co-cultures of HEK293 cells with rat hippocampal neurons, overexpression of SynCAM 1 in the HEK293 cells drives functional presynapse formation at the sites where axonal processes of hippocampal neurons contact them (Biederer et al., 2002; Fogel et al., 2007). We performed coculture experiments of hippocampal neurons with HEK293 cells expressing TMEM24-mCh in addition to SynCAM 1(363)-eGFP (see Figure 7A for diagram). In these cultures we observed strand-like TMEM24-positive ER/PM junctions at sites where neuronal processes made contacts with HEK293 cells, as assessed by anti-Tau or anti-synaptophysin immunofluorescence (Figure 7B). We also coplated neuroblastoma IMR32 cells expressing TMEM24 tagged with GFP at the endogenous locus with rat hippocampal neurons expressing the PM marker mCh-CAAX. Also in this system we found an enrichment of endogenous TMEM24 under neuronal processes (Figure 7C), showing that such localization occurs at physiological levels of expression of the proteins involved.

**Figure 7.**
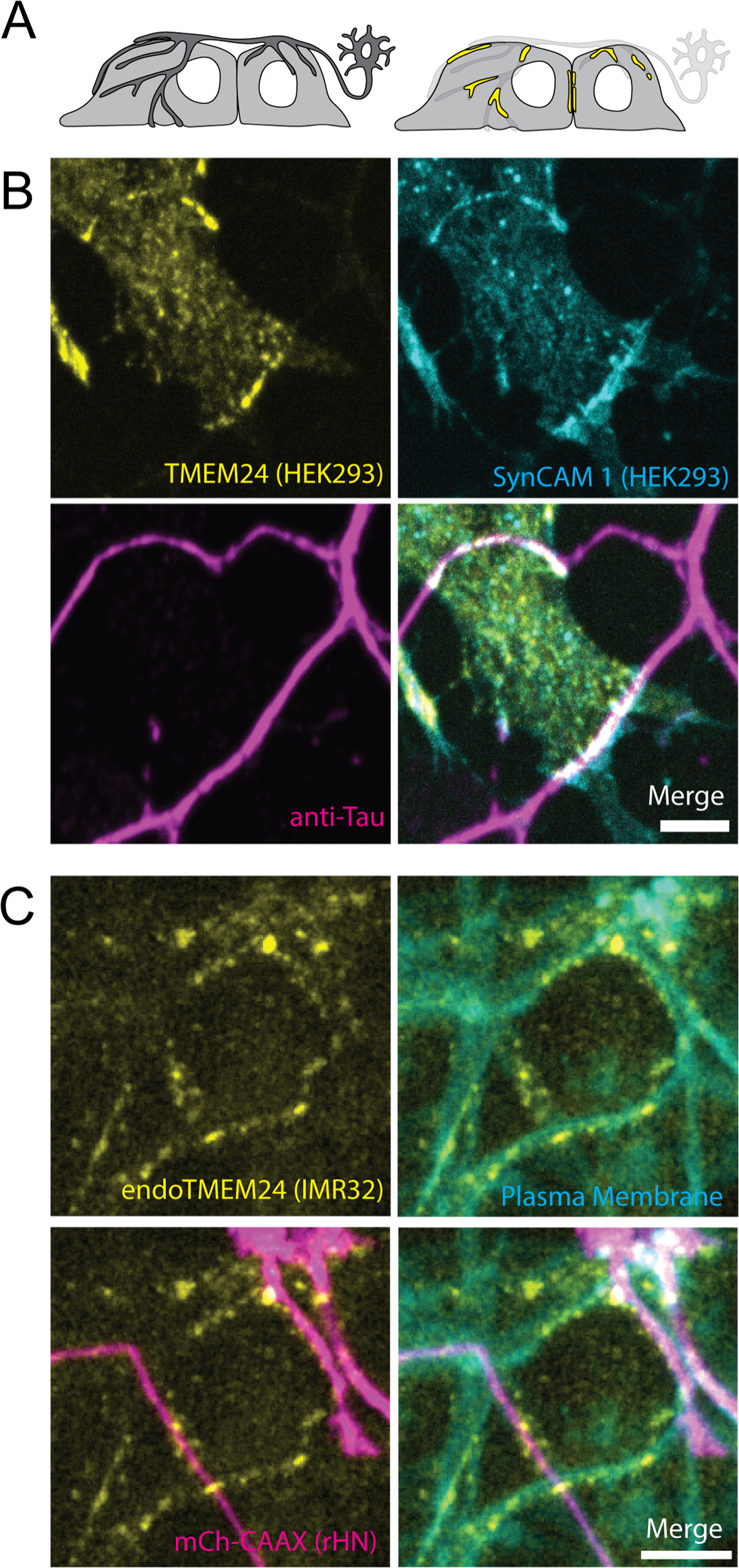
TMEM24 localizes to neurons contact sites. **(A)** Diagram of the coculturing system. **(B)** TMEM24-mCherry and SynCAM1(563)-eGFP expressed in HEK293 cells accumulate at contacts with axons of co-plated rat hippocampal neurons as revealed by anti-tau immunofluorescence. **(C)** Endogenous TMEM24 (endo-eGFP) in IMR32 cells accumulates at contacts with three neuronal processes when co-plated with rat hippocampal neurons expressing the PM marker mCherry-CAAX. Note that neurons are only sparsely transfected with mCherry-CAAX, so that at least some linear arrays of endo-eGFP spots not in register with mCherry-labeled axons may correspond to unlabeled axons. Scale bars = 5 μm

## DISCUSSION

The present study identifies a mechanism that couples cell-cell adhesion to the formation of ER/PM junctions with lipid transport properties. Our studies of TMEM24 suggest that this protein and its close paralogue C2CD2 are key players in this coupling. They do so through a mechanism that is distinct from the previously described property of both proteins to tether the ER to the PM via the binding of their C-terminal polybasic motif to the acidic cytosolic leaflet of the PM.

The accumulation of TMEM24 and C2CD2 at cell-cell interfaces is mediated by the interaction between an evolutionarily conserved small β-sheet motif within the predominantly disordered C-terminal region of TMEM24 and a β-sheet module in the CTD of the protein band 4.1 family. Based on AlphaFold multimer predictions these two β-sheets interlock with each other to form a single continuous chimeric β-sheet. Band 4.1 proteins bind a variety of plasma membrane proteins through their FERM domain. Thus, they can function as bridges between the ER and the PM. Importantly, as several interactors of band 4.1 proteins are cell adhesion molecules, the interaction of TMEM24 with band 4.1 provides an explanation for the concentration of TMEM24 at cell-cell contacts. Accordingly, our study shows that in adjacent HEK293 cells, tagged-TMEM24, band 4.1 and SynCAM 1 cocluster at symmetric ER/PM junctions. Interestingly, homotypic interactions between HEK293 cells mediated by complexes comprising band 4.1 proteins and SynCAM 1 which also comprise members of the MPP protein family have been described, all of which were hits in our screen for TMEM24 neighbors at sites of cell adhesion. However, it is quite possible that other cell adhesion proteins that bind band 4.1 besides SynCAM 1, (e.g., CD44, CADM4, β-integrins, and others) may contribute to the localization of protein 4.1, and thus TMEM24, at sites of cell-cell contacts. For example, in our cocultures of neurons with cells expressing tagged endogenous or exogenous TMEM24, cell adhesion proteins other than SynCAM 1, but that bind protein 4.1, may have contributed to the formation of ER/PM junctions at sites of cell adhesion.

As the presence of acidic phospholipids on the cytosolic leaflet is a general feature of the entire PM, the interaction of TMEM24 with protein 4.1 (via the β-sheet motif) and with the acid bilayer (via the polybasic motif) may synergize at sites of cell-cell contacts. The charge-based interactions of the polybasic motif may help strengthen the interaction of TMEM24 and C2CD2 with the PM mediated by 4.1 proteins and do so in a regulated way in the case of TMEM24.

Presence of ER/PM junction at specialized cell-adhesion sites has been observed in a variety of context in different tissues, implying the occurrence of mechanisms to coordinate extracellular interactions with the focal recruitment of the ER at these sites. For example, ER/PM junctions are present at cell-cell junctions along the basolateral surface of epithelial cells (Chung et al., 2022). Symmetrically arranged ER/PM junctions have been described at homotypic cell-cell contacts, including contacts between neuronal somata in the nervous system. Striking examples of asymmetric junctions where ER/PM cisterns are present in only one of the two participating cellular elements, are synapses where an ER cistern with a narrow lumen is closely apposed to the entire post-synaptic membrane such at synapses between C-fibers on motor neurons and between efferent axons and hair cells of the vestibular system (Deardorff et al., 2014; Smith and Sjöstrand, 1961). TMEM24 and C2CD2 could cooperate with other tethers to form some of these structures, as other mechanisms have also been described to generate ER/PM junctions at sites of cell adhesion. For example, the Kv2.1 potassium channel, which mediates ER/PM junctions by interacting with VAP, binds the cell adhesion PM protein AMIGO. There is vast literature on Kv2.1-positive ER/PM junctions being localized to areas of cell-cell contact, both at GABAergic synapses and at contact sites between neurons and astrocytes or microglia where the channel is thought to modulate neuronal activity, synaptic transmission, and response to ischemic insults (Cserép et al., 2020; Du et al., 1998; Misonou et al., 2008; Panzera et al., 2022).

The specific feature of TMEM24 and C2CD2 containing ER/PM junctions, however, is the presence of a lipid transport module in these proteins, implying that lipid transport is a process that occurs at these sites. Cell-cell contacts have an important role in intercellular signaling. Many of the membrane bound ligands and receptors that participate in mediating these cell interactions trigger intracellular signaling cascades which include phosphorylation of PI(4,5)P_2_ to PI(3,4,5)P_3_, PLC-mediated PI(4,5)P_2_ cleavage and other lipid metabolic reactions. Thus, it is plausible that TMEM24 and C2CD2 may participate in homeostatic responses to lipid perturbations mediated by such reactions. In this context it is of interest that at least some of the synapses with a post-synaptic cistern mentioned above, for example C-fibers synapses, are cholinergic with the presence post-synaptically of phospholipase C-coupled muscarinic receptors. Thus, cleavage of PI(4,5)P_2_ triggered by acetylcholine release at these sites to mediate some of its action requires lipid exchanges between the PM and the ER to replenish the depleted PI(4,5)P_2_ pool, a process in which TMEM24 and C2CD2 may participate (Lees et al., 2017; Xie et al., 2022).

Finally, proteins of the 4.1 family bind a plethora of cell surface proteins beyond proteins specialized for cell adhesion. Example of such ligands include AMPA receptors, NMDA receptors, mu-opioid receptors, and metabotropic glutamate receptors. Thus, although our present findings have emphasized an action of TMEM24 at sites of cell adhesion, the interaction of TMEM24 and C2CD2 with proteins of the band 4.1 family may function in ER/PM cross-talk in a variety of additional contexts.

## MATERIALS & METHODS

### Plasmids

TMEM24-eGFP, TMEM24-mCherry, TMEM24(1-414)-eGFP, TMEM24(1-630)-eGFP, TMEM24(1-666)-eGFP, TMEM24(5S→E)-eGFP, and C2CD2-eGFP constructs were previously reported (Lees et al., 2017; Sun et al., 2019).

Kv2.1-eGFP, Kv2.1loopBAD, and hBirA were kind gifts from Dr. M. Tamkun at Colorado State University. GFP-ESyt2 was described previously (Giordano et al., 2013). YFP-STIM1(D76A) was a gift from Dr. G. Voeltz (Addgene #186622). M1R was a gift from Dr. B. Hille.

TRAPγ was a kind gift from Dr. M. Mariappan at Yale University. TRAPγ-GFP was subsequently created by cloning TRAPγ into the EGFP-N1 vector using XhoI and HindIII cut sites. TRAPγ-TMEM24(1-414)-eGFP was generated using the TRAPγ-GFP and TMEM24(1-414)-GFP plasmids. TMEM24(ΔBS + 5S → E)-eGFP was generated using site-directed mutagenesis to remove a portion of the C-terminus of TMEM24(5S → E)-eGFP including the first 3 β-strands of the β-sheet band 4.1 interacting domain (Quik-Change II XL; Agilent Technologies).TMEM24(5S → E)-eGFP which has been previously described (Sun et al., 2019). eGFP-Sec61β-APEX2 was cloned using eGFP-Sec61β and eGFP-APEX2 and XhoI and XmaI cut sites. TMEM24-APEX2-eGFP was cloned using Tom20-APEX2-eGFP and TMEM24-eGFP and XhoI and XmaI cut sites. TMEM24(1-414)-APEX2-GFP, TMEM24(1-630)-APEX2-GFP, TMEM24(1-666)-APEX2-GFP were generated from TMEM(1-414)-GFP, TMEM24(1-630)-GFP, and TMEM24(1-666)-GFP respectively using TMEM24-APEX2-eGFP and standard restriction enzyme cloning techniques.

mCherry-4.1G, mCherry-4.1G(ΔCTD), and Mem-mCherry-4.1G(CTD) were generous gifts from the I. Cheeseman lab (Addgene plasmids #46361, #46360, and #46362, respectively). eGFP-4.1R was generated by using pTK81_GFP-4.1R (Again a gift from I. Cheeseman, Addgene #46352) and cloning into CMV-eGFP-C1 vector using XhoI and SacII cut sites.

SynCAM 1(363)-GFP has been previously described (Fogel et al., 2007)

### Cell Culture & Transfection

HEK293 cells were cultured at 37°C and 5% CO_2_ in DMEM (Gibco, cat 11965-092) + 10% FBS. IMR32 neuroblastoma cells with TMEM24 endogenously tagged with GFP was accomplished via CRISPR/Cas9 by the company PNAbio and were previously described (Sun et al., 2019). IMR32 cells were cultured at 37°C and 5% CO_2_ in DMEM (Gibco, cat 11965-092), 15% FBS, 1 mM sodium pyruvate (Gibco, cat 11360-070), and 2 mM glutaMAX (Gibco, cat 35050-061). For differentiation, IMR32 cells were cultured in media supplemented with 2.5 µM bromo-deoxyuridine. Differentiation medium was replaced by 50% every 2-3 days and cells were used for experiments after 14-21 days of differentiation when clear neuronal-like processes could be seen by simple light microscopy. HEK293 cells, IMR32 cells, and differentiated IMR32 cells were transfected using lipofectamine 2000 (Invitrogen, cat 52887) and used for experiments the following day. For HEK293-HEK293 co-culture experiments, populations of cells were transfected separately, allowed to express the protein of interest overnight, and then plated together on Mattek dished the following morning. Microscopy was performed later that afternoon/evening after the cells had several hours to settle and adhere to the new dish. All cells were checked for mycoplasma contamination monthly. For HEK293-rat hippocampal co-culture experiments we used a previously published protocol (Biederer and Scheiffele, 2007). Briefly, rat hippocampal neurons were collected from postnatal day 0 or 1 animals and grown in culture until day in vitro 7-10. Simultaneously, HEK293 cells were transfected with the protein of interest (TMEM24-mCherry, mCherry-JPH4, and/or SynCAM 1(363)-GFP; see individual experiment for construct details) and seeded atop the neuronal culture the following day. 24-48 hours after combining the cultures the dishes were fixed in DPBS + 4% formaldehyde for 15 min. Cell membranes were permeabilized with 0.5% CHAPS in DPBS and blocked before being labeled with either rabbit anti-synaptophysin (Synaptic Systems, 101 002) or mouse anti-tau (Cell Signaling, 4019S) primary antibodies and either donkey anti-rabbit AlexaFluor 647 (Invitrogen, A31573) or goat anti-mouse AlexaFluor 680 (Invitrogen, A21058) secondary antibodies. All work with rats was performed in accordance with both Yale institutional and federal guidelines.

### Microscopy

Imaging was performed using an Andor Dragonfly spinning-disk confocal imaging system with a Zyla CMOS camera and 60x plan apochromat objective (63x, 1.4 NA, oil). For live cell experiments, cell media was replaced with HEPES-buffered live-cell imaging solution (Invitrogen, cat A14291DJ) and imaging was performed at 37° C and 5% CO_2_. Labeling with Cellbrite and Halo dyes and wash steps were done in imaging saline immediately prior to imaging. CellBrite (Biotium, cat 30108) was used at 1:1000 for 7-10 minutes before washing with imaging saline. Janelia JF646 HaloTag was used at 1:1000 for 10 minutes before wash.

### Oxotremorine-M Treatment

For the Oxotremorine-M dissociation of TMEM24, cells were transfected with TMEM24-mCh, M1R(unlabeled) and stained using a far-red Cellbrite PM dye. During image acquisition, after a baseline was established, Oxo-M dissolved in imaging saline was added to the dish to a final concentration of 10 μM while TMEM24 localization was recorded.

### Image Processing and Analysis

All microscopy image processing was performed using ImageJ software. Images were pseudocolored, cropped, and adjusted for contrast and brightness. Background subtraction, filtering, and noise removal, where necessary, were performed equally across all channels to generate clearer pictures for publication. Analysis was performed on either unprocessed images or images that had undergone equal background subtraction across channels but no further processing. For normalization, averages of pretreatment values were used as a baseline.

### APEX2 Proteomic Analysis

Our APEX2 protocol is based on the published protocol of the Ting laboratory (Hung et al., 2016). Briefly, HEK293 cells were transfected with the indicated plasmids and allowed to express the proteins over a 24-hour period. Cells were then incubated with 500 μM biotin tyramide (Iris Biotech, cat 41994-02-9) in DMEM + 10% FBS for 1 hr, treated with 1 mM H_2_O_2_ for 1 min to induce the APEX reaction, treated with quenching solution (10 mM sodium ascorbate, 5 mM Trolox and 10 mM sodium azide, in DPBS) to end radical formation, washed 3x in PBS and either fixed in DPBS + 4% formaldehyde for 15 min (if the cells were to be imaged) or collected in DPBS + protease inhibitor (for affinity purification). Cells that were imaged were labeled using CF640-conjugated streptavidin to label biotinylated proteins (CF640R Streptavidin, Biotium cat 29041). Affinity purification was performed using streptavidin-conjugated magnetic beads (Pierce, REF 88817) utilizing the steps and buffers of the previously referenced protocol [13]. The affinity purified samples were then either submitted for mass spectrometry analysis or subjected to gel electrophoresis to generate the western blot figures found in the manuscript. Membranes were probed with mouse anti-GFP (Sigma, G6539) then labeled with LiCor IRDye 680LT Goat α-Mouse (926-68020) and LiCor IRDye 680RD Streptavidin (926-68079). Membranes were imaged on a LiCor Odyssey Classic system. LC MS/MS with label free quantitation was performed on purified samples in triplicate after a precipitation step to reduce detergent and free biotin concentration. A threshold of 2+ unique peptides identified was used and proteins that did not satisfy this requirement were not included in the analysis.

### Mass Spectrometry Sample Preparation

APEX eluted protein solutions were submitted to the Keck MS & Proteomics Resource at the Yale School of Medicine for mass spectrometry analyses. Proteins were extracted utilizing a cold acetone protein precipitation method. Briefly, 400uL cold acetone (−20°C) was added to the ∼90uL eluted protein samples and vortexed. The precipitation was allowed to continue overnight in a −20°C freezer; then the mixture was immediately centrifuged at 14.6K at 4°C for ten minutes. The pellet was then air dried (but not to completeness); then reconstituted in 20uL 0.1% Rapigest (Waters Inc., Milford, MA) containing ammonium bicarbonate (ABC). The proteins were then reduced with 2uL 45 mM dithiothreitol (DTT) at 37°C for 30 minutes and cooled to room temperature. Alkylation with 2 uL 100 mM iodoacetamide at room temperature for 30 minutes in the dark. 2uL of trypsin (1:5 of a 0.5ug/uL trypsin in H_2_O) was then added and incubated at 37°C overnight (e.g., 16 hours). Digest samples were then quenched (and Rapigest was crashed out) with 1.3uL 20%TFA in 37°C for 45 minutes. Supernatant containing the peptides for analyses was then moved to a new Eppendorf tube. An aliquot was taken, and concentration measured via Nanodrop, and diluted to 0.05 µg/µl with 0.1% TFA. 1:10 dilution of 10X Pierce Retention Time Calibration Mixture (Cat#88321) was added to each sample prior to injecting on the UPLC Q-Exactive Plus, to check for retention time variability the normalization during LFQ data analysis.

### Mass Spectrometry

Peptides were analyzed by LC–MS/MS using either a Q-Exactive Plus mass spectrometer equipped with a Waters nanoACQUITY ultra-performance liquid chromatography (UPLC) system using a Waters Symmetry C18 180 mm by 20 mm trap column and a 1.7 mm (75 mm inner diameter by 250 mm) nanoACQUITY UPLC column (35°C) for peptide separation. Trapping was done at 5µl/min, 99% Buffer A (100% water, 0.1% formic acid) for 3 min. Peptide separation was performed at 300 nl/min with a linear gradient that would reach 5% Buffer B (100% CH_3_CN, 0.075% formic acid) at 2 min, 25% B at 140 min, and 40% B at 165 min, and 90% B at 170 min for 10 min; then dropped down to 3% B at 182 min for 5 min. For the LCMS/MS data dependent acquisition on the Q-Exactive Plus mass spectrometer, High-energy Collisional Dissociation (HCD) MS/MS spectra were filtered by dynamic exclusion (20 seconds) and acquired for the Top 20 peaks with charge states 2-6 with m/z isolation window of 1.7. All MS (Profile) and MS/MS (centroid) peaks were detected in the Orbitrap.

### MS Data Analysis

Mass spectral data were processed using Progenesis QI (Waters Inc., Milford, MA; v.4.2). Analysis method is described elsewhere by (Torregrossa et al., 2018). Briefly, Peaks were picked in Progenesis QI, and LC MS/MS mascot generic file (.mgf) were exported for protein search with inhouse MASCOT Search engine. Protein searches were conducted against the Homo sapiens SWISSProt protein database using Mascot Search Engine (Matrix Science, LLC, Boston, MA; v. 2.6.0). Mascot search parameters included: parent peptide ion tolerance of 10.0 ppm, peptide fragment ion mass tolerance of 0.020 Da, strict trypsin fragments (enzyme cleavage after the C terminus of K or R, but not if it is followed by P), variable modification of phospho (S, T, Y, and H), oxidation (M), and carboxyamidomethyl (C).

The mass spectrometry proteomics data have been deposited to the ProteomeXchange Consortium via the PRIDE (Perez-Riverol et al., 2022) partner repository with the dataset identifier PXD047112.

## Acknowledgements

We thank A. Guillén-Samander and J. H. Park for technical and cloning assistance and A. Paquette for preparing neuronal cultures. This work was supported in part by the NIH grants R01 NS36251 and DA018343 (to P.D.C.) and R01 DA018928 (to T.B.) and the Kavli Institute for Neuroscience. We also thank the MS & Proteomics Resource at Yale University for providing the necessary mass spectrometers and biotechnology tools funded in part by the Yale School of Medicine and by the Office of The Director, National Institutes of Health (S10OD02365101A1, S10OD019967, and S10OD018034). The funders had no role in study design, data collection and analysis, decision to publish, or preparation of the manuscript.

## Author Contributions

B. Johnson and P. De Camilli conceptualized the project. T. Biederer provided critical advice and tools. B. Johnson and M. Iuliano together performed the HEK293-neuronal coculture experiments and analysis. LC MS/MS was performed by T.K. Lam and the Yale MS & Proteomics Resource Center after APEX sample preparation by B. Johnson. All other experiments and analysis were performed by B. Johnson. B. Johnson and P. De Camilli wrote the original draft of the manuscript which was subsequently reviewed and edited by all authors.

---

**Supplemental Figure 1.**
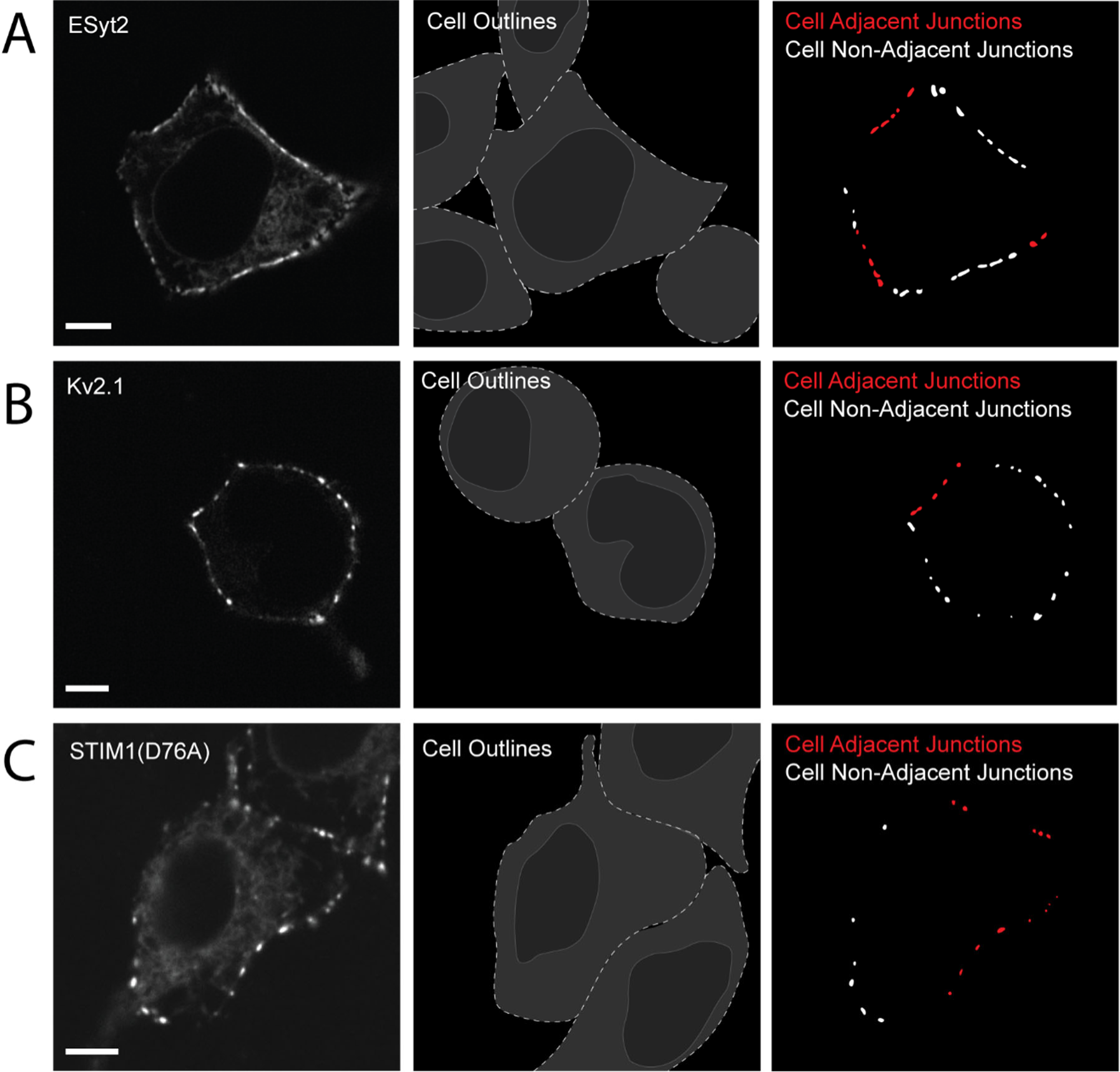
Several ER/PM junction tethers tested other than TMEM24 and C2CD2 display no preference for cell-adjacent ER/PM junctions. ER/PM junctions positive for exogenously expressed E-Syt2-eGFP **(A)**, Kv2.1-eGFP **(B)** or YFP-STIM1(D76A)**(C)** in HEK293 cells show no differences in size depending on cell adjacency. Quantification of this data can be found in Figure 1B of the main text. Scale bars = 5 μm.

**Supplemental Figure 2.**
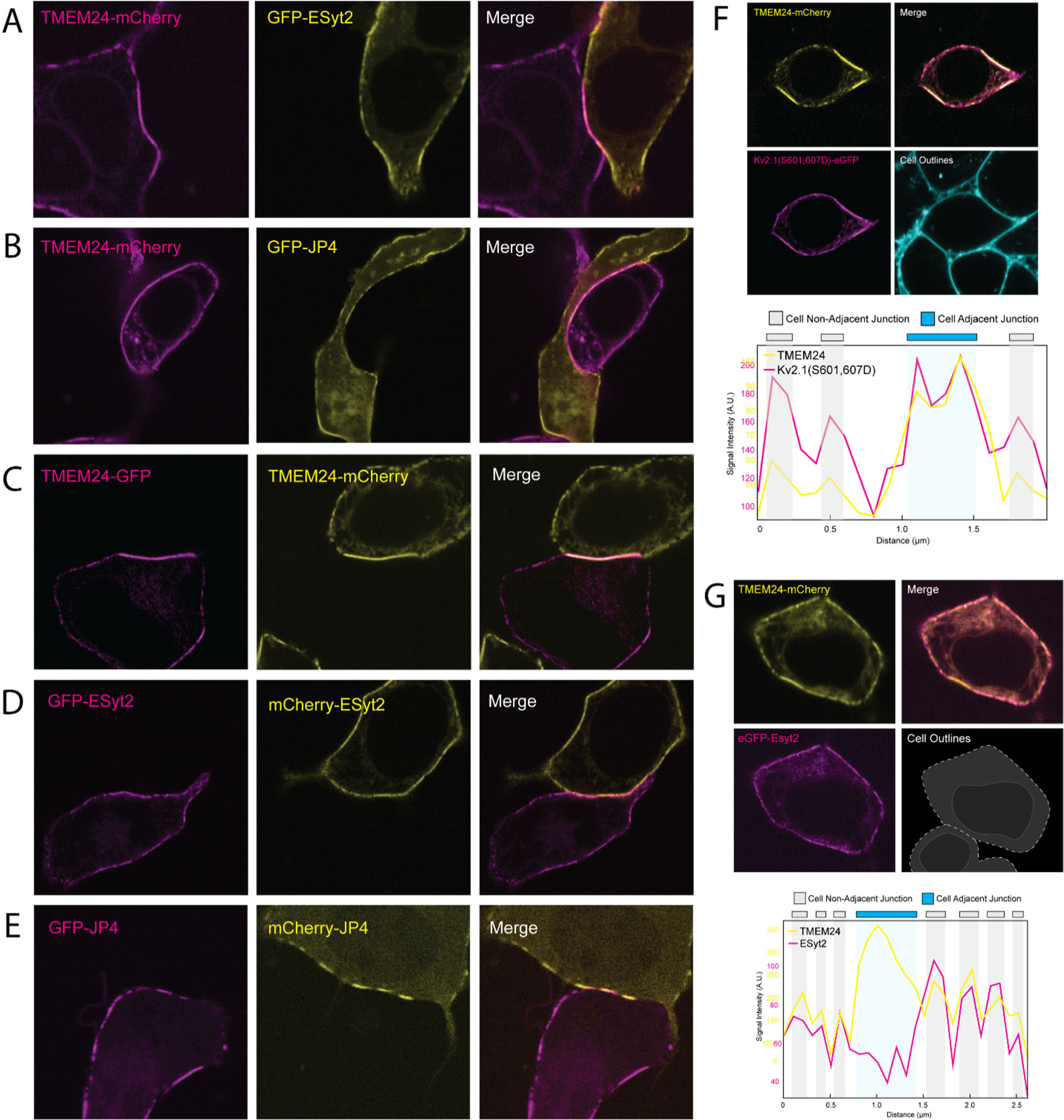
ER/PM Junction Tether Behavior at the Cell-Cell Interface. HEK293 cells. TMEM24 expressed in one cell generates an enlarged ER/PM junction that is not mirrored by an EGFP-ESyt2-positive **(A)** or an JPH4-(EGFP)-positive **(B)** junction in an adjacent cell. **(C)** The accumulation of TMEM24-mCherry expressed in one cell at a cell-adjacent ER/PM junction is mirrored by the accumulation of TMEM24-GFP expressed in an adjacent cell (see also Figure 1C). **(D)** ER/PM junctions induced by mCherry-ESyt2 and GFP-ESyt2-expressed in two adjacent cells respectively, do not mirror one another across the cell-cell interface. **(E)** mCherry-JPH4 and GFP-JPH4 do not robustly mirror one another across the cell-cell interface although junctions could be found that seemed to be symmetrically opposed. **(F** and **G)** A Kv2.1-GFP mutant constitutively bound to VAP **(F)**, but not EGFP-E-Syt2 **(G)**, colocalizes with TMEM24 at cell-adjacent ER/PM junctions. At non cell-adjacent junctions, TMEM24 colocalizes with both proteins. Quantification of line scans drawn around the cell periphery are shown below the fluorescence images.

**Supplemental Figure 3.**
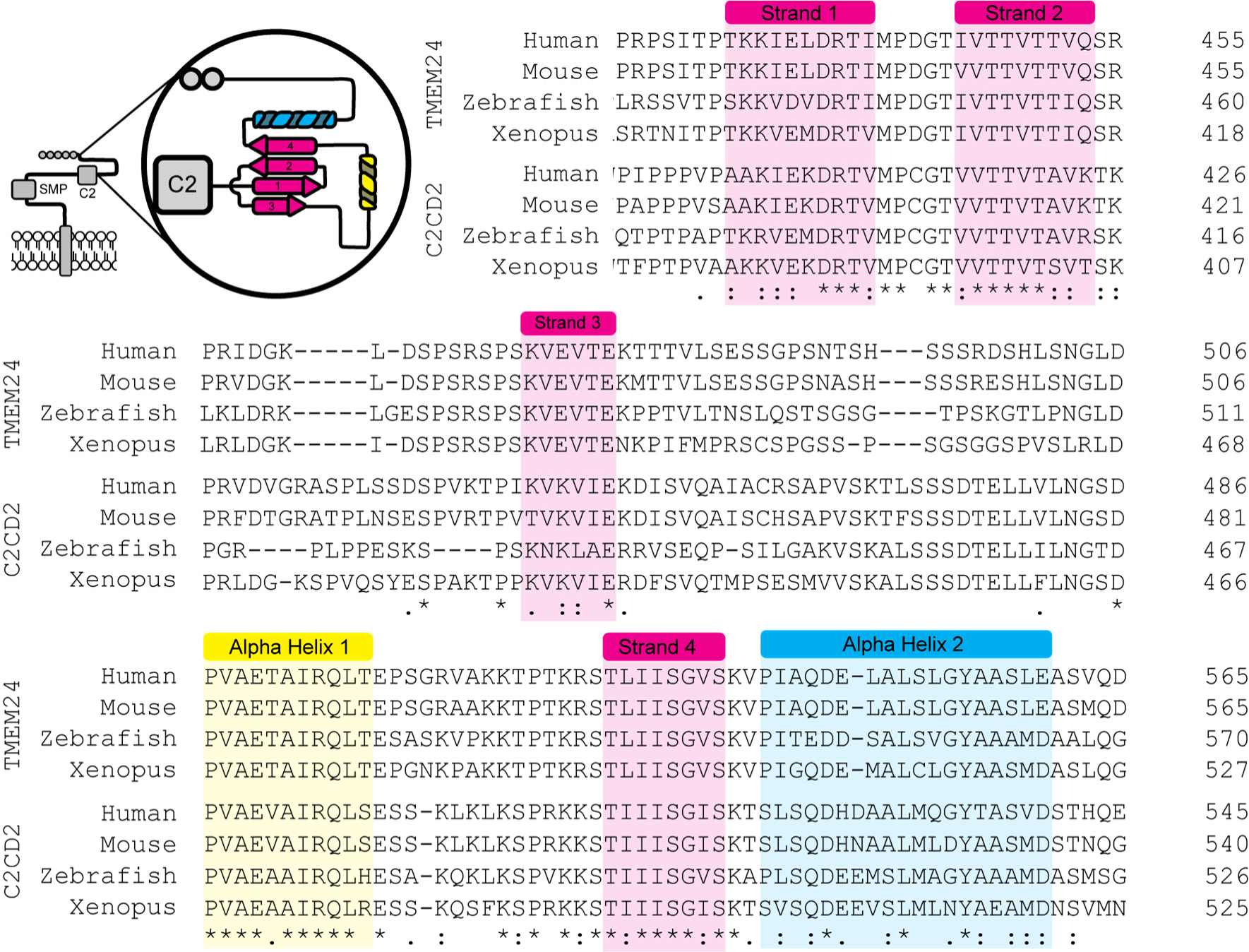
Sequence alignments of portions of the TMEM24 and C2CD2 C-termini across species demonstrating conservation of the *α*-helixes and *β*-strands. The cartoon at top left shows a schematic view of a TMEM24 monomer, with an enlarged view of the Alphafold predicted structural motifs (colored) within its 414-630 amino acid region.

**Supplemental Figure 4.**
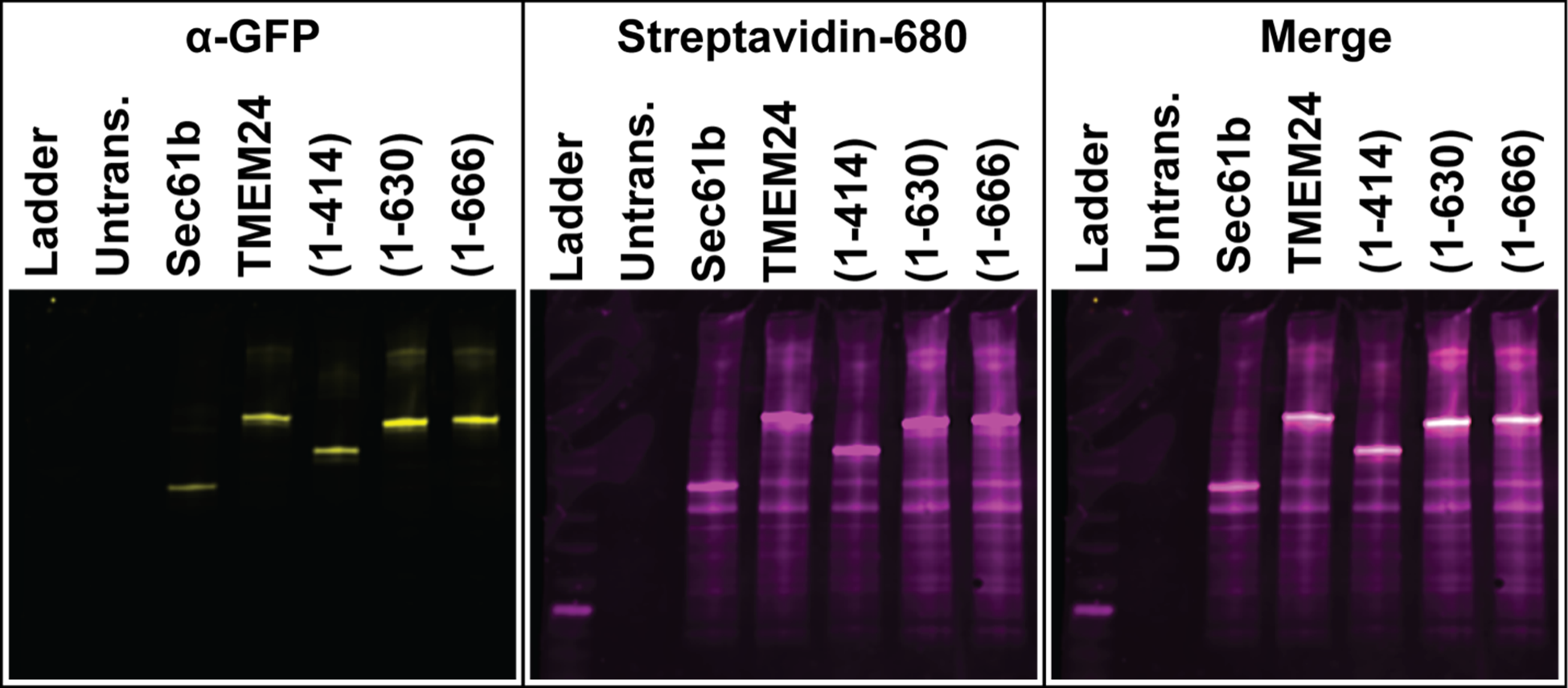
Streptavidin affinity-purification of cell extracts expressing the constructs indicated and homogenized after the APEX2 reaction. Anti-GFP Western blots and streptavidin overlay of material affinity-purified on streptavidin bead. Numbers in parenthesis indicate amino acid boundaries of TMEM24 fragments used. Note the high degree of self-biotinylation for each construct.

**Supplemental Figure 5.**
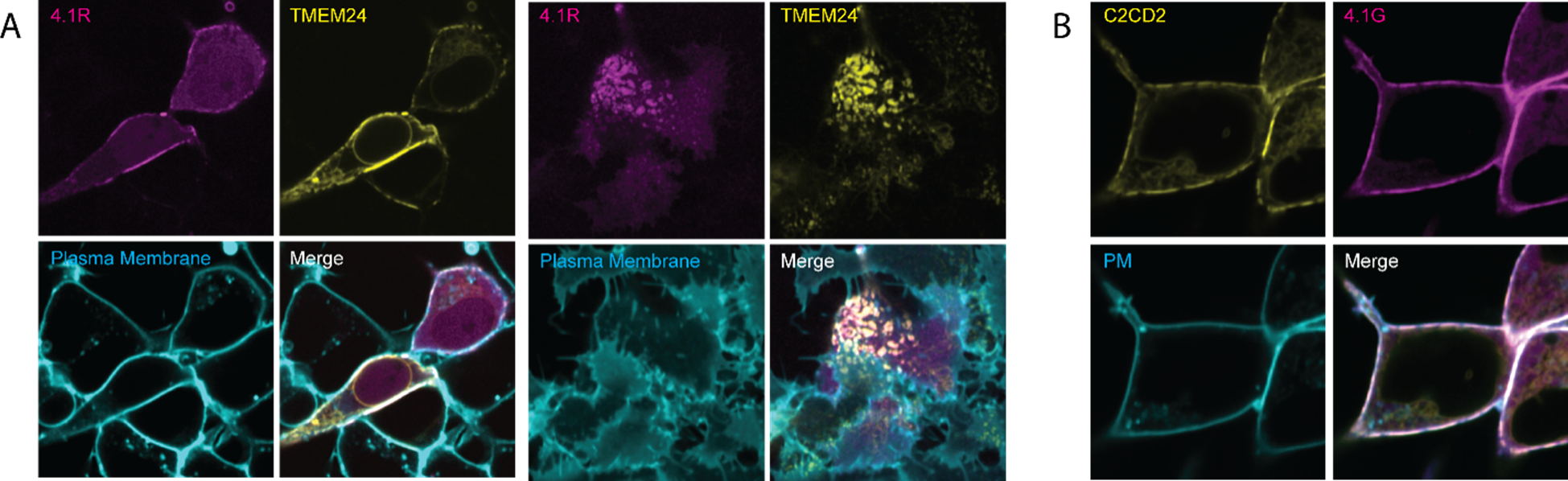
The TMEM24-Band 4.1 partnership is observed also with their paralogues. **(A)** Colocalization of TMEM24-mCherry with band 4.1R-GFP at ER/PM junctions of HEK293 cells as seen in both a mid-cell confocal z-slice and at the basal surface. **(B)** Colocalization of C2CD2-eGFP with mCherry-4.1G.

